# Snap Back to Reality: Comparing Real-World Spatial Memory in the “Wild” vs. in the Lab

**DOI:** 10.64898/2026.03.25.714295

**Authors:** Derek J. Huffman, Paisley J. Annes, Chandrachud Gowda, Lior Colina

**Author notes:** Correspondence should be addressed to Derek J. Huffman, Department of Psychology, Colby College, Waterville, ME 04901, USA.

## Abstract

We thank the Department of Psychology at Colby College for supporting this work.

Derek J. Huffman played a primary role in conceptualization, funding acquisition, project administration, methodology, resources, supervision, writing–initial draft, and writing–review and editing; and a supporting role in formal analysis, software, validation, and visualization. Paisley J. Annes played a lead role in validation and visualization; and a supporting role in conceptualization, data curation, formal analysis, methodology, software, writing–initial draft, and writing–review and editing. Chandrachud Gowda played a primary role in Investigation; and a supporting role in conceptualization, data curation, formal analysis, methodology, project administration, visualization, and software. Lior Colina played a supporting role in conceptualization, methodology, and software.

Memory is most commonly studied using lab-based measures; however, spatial memory for large-scale environments, such as cities, typically takes extended time and motivation to learn, thus raising questions about the ecological validity of findings from traditional lab-based paradigms (e.g., information that is learned over a short timescale and with no real-world relevance). We leveraged technological breakthroughs to create a custom smartphone application to compare situated spatial memory with lab-based measures. Specifically, we compared performance in healthy young adults across two established lab-based tasks, judgments of relative direction (JRD) and map drawing, and our novel app-based, in situ pointing task for the same familiar, large-scale, real-world environment (the participants’ college campus). We observed a strong correlation between JRD performance and map-drawing performance. App-based pointing showed lower error and reduced inter-individual variability relative to JRD performance, but weak correlations with lab-based measures. We also developed a novel analytical technique in which we transformed the app-based pointing into a relational, JRD-like metric, and we observed strong correlations with the lab-based tasks. Moreover, we observed correlated patterns of errors across all tasks, thus suggesting that these tasks tap into partially overlapping cognitive representations. Our results provide a pivotal advancement to our understanding of both shared and unique variance across spatial memory paradigms and support the use and further development of mobile navigation tools as complements to lab-based assessments for studying spatial cognition and differences across populations, including patients with Alzheimer’s disease who often report getting disoriented even within familiar spatial environments.

## 1. Introduction

Spatial memory supports everyday functioning. For example, memory for familiar environments supports independent living across the lifespan (Wolbers & Hegarty, 2010). The dominant approach for studying spatial memory includes asking participants to learn new environments and then asking them to make spatial memory judgments about those environments. Since the advent of video game engines and more immersive VR technology, many lab-based studies have also employed virtual environments and tasks (Coughlan et al., 2019; Coutrot et al., 2018, 2022; Richardson et al., 1999; Spiers et al., 2023; Waller et al., 2004; Weisberg et al., 2014). Moreover, researchers have developed several lab-based tasks to study spatial memory, including judgments of relative direction (JRD) and Map-Drawing, and these types of tasks have played an important role in characterizing spatial memory (Hegarty et al., 2006; Labate et al., 2014; Mou et al., 2004, 2007; Mou & McNamara, 2002; Rieser, 1989; Shelton & McNamara, 1997, 2001a, 2001b; Weisberg et al., 2014), especially related to abstract, holistic memory for the configuration of landmarks within large-scale environments (Ekstrom et al., 2017; Huffman et al., 2026; Huffman & Ekstrom, 2019; Marchette et al., 2011; McNamara et al., 2003; Nakaya, 1997). These tasks have demonstrated that spatial cognition relies on structured internal representations that support flexible reasoning about locations and directions. Despite the theoretical importance of laboratory-based paradigms, these approaches may be limited in ecological validity and scalability, as they typically require abstract judgments that are spatially and contextually removed from the environments that they are intended to measure (Diersch & Wolbers, 2019; Taube et al., 2013; Wang & Spelke, 2002), thus the comparison of lab-based tasks with tasks in which the navigator is situated in the environment can provide important insight into whether these tasks recruit overlapping cognitive representations.

One approach for testing whether tasks tap into common cognitive representations is to compare correlations across spatial memory tasks both between and within individuals, and the observation of correlations between tasks is often taken as evidence that tasks tap into partially overlapping cognitive representations (e.g., Bryant, 1984; Coughlan et al., 2019; Coutrot et al., 2019; Hegarty et al., 2006; Huffman & Ekstrom, 2019); however, important questions remain about how well such findings would relate to the comparison of spatial memory performance in the laboratory vs. in situ. For example, previous studies have found correlations between how well people perform on various spatial memory tasks for experiments in which participants learn a new spatial environment, including the JRD Task and map drawing/model building tasks (e.g., Huffman & Ekstrom, 2019). Moreover, past studies have argued that the best tests for analyzing the similarity of cognitive representations between spatial memory tasks include both correlations between tasks (i.e., individual differences) and between patterns of errors for equivalent trial-wise measures (Bryant, 1984; Huffman & Ekstrom, 2019). In line with these ideas, recent research has suggested that the patterns of errors are correlated for lab-based measures of spatial memory (e.g., the JRD Task and the Map Drawing Task) both for a virtual environment (Huffman & Ekstrom, 2019) and familiar large-scale, real-world environments (Huffman et al., 2026). Here, we aim to determine whether such findings replicate and extend to the comparison between the laboratory vs. in situ behavioral performance.

Recent advances in mobile and digital technologies offer exciting new opportunities to assess spatial navigation in more naturalistic contexts; however, relatively few studies have compared performance for in situ tasks vs. the laboratory. On the one hand, some recent studies have found a correlation between learning of a new virtual environment and learning a new real-world environment (Coutrot et al., 2019; Puthusseryppady et al., 2022). On the other hand, research on active spatial learning suggests that physically engaging with environments can enhance spatial memory beyond what is captured by desktop-based virtual environments, passive navigation, or purely abstract and disembodied tasks (Chance et al., 1998; Chrastil & Warren, 2012, 2013, 2015; Diersch & Wolbers, 2019; Klatzky et al., 1998; Ruddle et al., 2011; Ruddle & Lessels, 2006; Wang & Spelke, 2000), with some theories placing key importance on body-based cues for both spatial learning and memory performance (e.g., Taube et al., 2013; Wang & Spelke, 2002). Moreover, previous research has suggested that there is a partial dissociation between JRD performance and performance of in situ pointing tasks in which participants are oriented within their environment (e.g., Waller & Hodgson, 2006; Zhang et al., 2014). Altogether, these findings raise two possible hypotheses for the comparison of human spatial memory performance in the lab vs. the real world. The first hypothesis predicts that spatial memory performance will be correlated between the lab and app-based measures; i.e., these tasks tap into a common underlying representation. The second hypothesis predicts that spatial memory performance will exhibit weak or null correlations between the lab and app-based measures; e.g., due to possible dissociations in cognitive processes, task demands, or access to body-based cues or other orienting cues. Here, to compare between these competing hypotheses, we developed and evaluated a novel app-based, in situ navigation measure alongside established lab-based JRD and map reconstruction tasks for memory for a familiar, large-scale environment: the participants’ college campus. We argue that both of the competing hypotheses are interesting and would provide important insight into theories of spatial memory, thus we employed a fully Bayesian analysis to determine which hypothesis was better supported by the data (Wagenmakers, 2007). We further argue that using a familiar, large-scale spatial environment is important for comparing these hypotheses because such environments are common in our everyday lives and can take extensive time and motivation to develop. Moreover, as we argue in the Discussion, it is possible that subtle signs of disorientation in familiar environments may be apparent in preclinical Alzheimer’s disease vs. absent in healthy aging.

To foreshadow our results, we found evidence of strong correlations within the lab-based tasks, thus supporting previous studies. For the comparison between our app-based, in situ pointing task and the laboratory-based tasks, we found evidence of both shared and unique variance, depending on the method of analysis. Specifically, when we compared the raw performance across the app-based task and the lab-based tasks, we observed very weak correlations but when we transformed the app-based pointing into a JRD-like metric, we then observed strong correlations in performance. Moreover, we observed strong correlations between the patterns of errors across all tasks, thus suggesting they tap into at least partially overlapping cognitive representations. In the Discussion, we highlight the importance of our findings, including a discussion of how our results provide a foundation for some initial ideas for further development of real-world spatial memory tasks for detecting preclinical memory changes in people at risk for Alzheimer’s disease, who often get disoriented even in familiar environments.

## 2. Method

### 2.1. Participants

We recruited 67 participants (41 female, 25 male, 1 non-binary) aged (18–22; 1 not reported) through the SONA recruitment pool, word of mouth, and school-sponsored advertisements. Participants received course research participation credit. All participants provided informed consent prior to participation, and all procedures were approved by the Institutional Review Board of Colby College.

We excluded participants from our analysis if they failed to complete both sessions, if their JRD performance or app-based pointing performance was not better than chance (as determined by a permutation procedure; see 2.3.3.; Huffman & Ekstrom, 2019), or if their map-drawing performance was not better than chance (as determined by the results from the bidimensional regression approach; see 2.3.3.; Carbon, 2013; Friedman & Kohler, 2003; Tobler, 1994). We excluded participants that performed at chance because 1) we did not want outliers to influence the results of any correlation analyses, 2) we assumed that if participants performed at chance, then they only contributed noise to our analysis (e.g., especially given the correlational nature of our analyses; Huffman & Ekstrom, 2019), 3) if participants performed at chance, then we cannot ensure that they put any effort into their performance in our experiment (in which case they would have chance performance across all tasks, thus artificially guaranteeing a correlation between our task measures). After applying the exclusion criteria, the final analyzed sample consisted of 58 participants.

Our a priori minimum sample size was to include at least 50 participants given the correlational nature of our main analyses (e.g., previous studies found strong correlations between individual differences and patterns of errors in performance on the JRD Task and the Map Drawing Task with 26 participants; Huffman & Ekstrom, 2019). Additionally, we employed a fully Bayesian analysis here (see 2.3.1.), thus we aimed to ensure that Bayes factors were greater than 3 in support of either the alternative or the null for the correlation between the JRD Task and the Map Drawing Task (both for the individual differences analysis and the partial correlation analysis; note that Bayesian statistics differ from frequentist statistics with respect to continuing data collection until a Bayes factor criterion is met, which would be deeply problematic in a frequentist approach; see 2.2. and 2.3.). We completed data collection of the present sample at the end of a semester of SONA participation. Upon processing and analyzing the data, we found that we met our criteria for the sample: we met both the target sample size (i.e., at least 50 participants) and the Bayes factors for the comparisons between the JRD Task and the Map Drawing Task (both the correlations and the partial correlations) were greater than 3 (in fact, all were much greater than 100). We did not preregister our study, but we based our analyses on previous studies (Huffman et al., 2026; Huffman & Ekstrom, 2019).

### 2.2. Task and Procedure

Here, we examined spatial memory for a familiar real-world environment: the Colby College campus. The study consisted of two sessions conducted within one week of each other.

#### 2.2.1. Session One: Computer-Based Tasks

In the Session One, we designed a lab-based task within Unity 3D that was built on top of the Landmarks package (Huffman et al., 2026; Starrett et al., 2021). Participants completed this session remotely in a full-screen application on a Windows computer. All participants performed this session first, rather than in a counterbalanced manner, because we did not want the participants to ever experience the real-world pointing angles prior to performing the JRD task (i.e., because then they could potentially use their real-world memory from those trials to help inform their performance on the JRD task).

##### Campus Familiarity Task

Participants first rated their familiarity with the location of all of the campus buildings using a 6-point Likert scale. We wrote code that selected the 10 campus buildings with the highest familiarity ratings, which we used for all of our spatial memory tasks.

##### Computer-Based Judgments of Relative Direction (JRD) Task

Participants then completed 100 trials of the JRD task, in which they estimated the direction between pairs of these 10 most familiar campus landmarks. For each trial, we asked participants questions with the following format: “Imagine that you are standing at A, facing B. Please point to C.” We randomly generated 100 JRD trials for each participant from the broader set of 720 possible questions for their 10 most familiar campus landmarks (i.e., 10 x 9 x 8).

We calculated the absolute angular error between the Response angles obtained during JRD trials along with the Answer angles (i.e., the ground truth or correct angles), which we calculated using the `bearing` function from the geosphere package in R and custom-written code based on the “arrival coordinates” from the app-based navigation task. We calculated the angular difference between the “pointing” angle (i.e., the bearing between the standing location and the pointing location) and the “facing” angle (i.e., the bearing between the standing location and the facing location). We then constrained these angles to be between 0 and 360 degrees (i.e., by adding 360 to any angles that were less than 0). We then calculated the absolute angular error between each participants response and the answer for each trail. Note that we used the participants’ “arrival” coordinates from each target location from the Navigation Task to calculate the answer angles (see 2.2.2.). We used the median absolute angular error as the primary performance measure to reduce the influence of outliers (i.e., the median is less influenced by outliers than the mean and here we had a relatively large number of trials, 100; for a similar approach see: Huffman et al., 2026; Huffman & Ekstrom, 2019).

##### Computer-Based Map Drawing Task

Participants next completed a Map Drawing Task for the same 10 landmarks. At the start of the task, circles with names above them were placed in a randomized order on the left-hand side of the screen. Participants used the mouse to select each corresponding landmark name (by clicking the circle) and move it to their desired location. Participants were required to move each one of the landmarks to be able to complete the map task (i.e., the task would not allow them to submit their response until they had moved each landmark from its initial starting location). The Map Drawing Task provided a measure of the participants’ overall understanding of the relative locations of the landmarks.

We analyzed the Map Drawing Task data via bidimensional regression (Carbon, 2013; Friedman & Kohler, 2003; Tobler, 1994). We used the Fisher r-to-z transformed (i.e., the inverse hyperbolic tangent function) model fit score from the Euclidean model (i.e., allowing for translation, rotation, and uniform scaling) as our primary measure for comparison of performance with the other tasks (as in previous research; e.g., Huffman & Ekstrom, 2019).

We generated a Map-Drawing-prediction of JRD performance by calculating the “heading” and “pointing” angles from the participants’ maps for each trial of the JRD task and then we calculated the (signed) angular difference between these angles (see Figure 1). We compared patterns of errors by employing a within-participant partial correlation approach on the JRD angles, the Map-Drawing-prediction of the JRD angles, and the Answer angles (see 2.3.1.).

**Figure 1:**
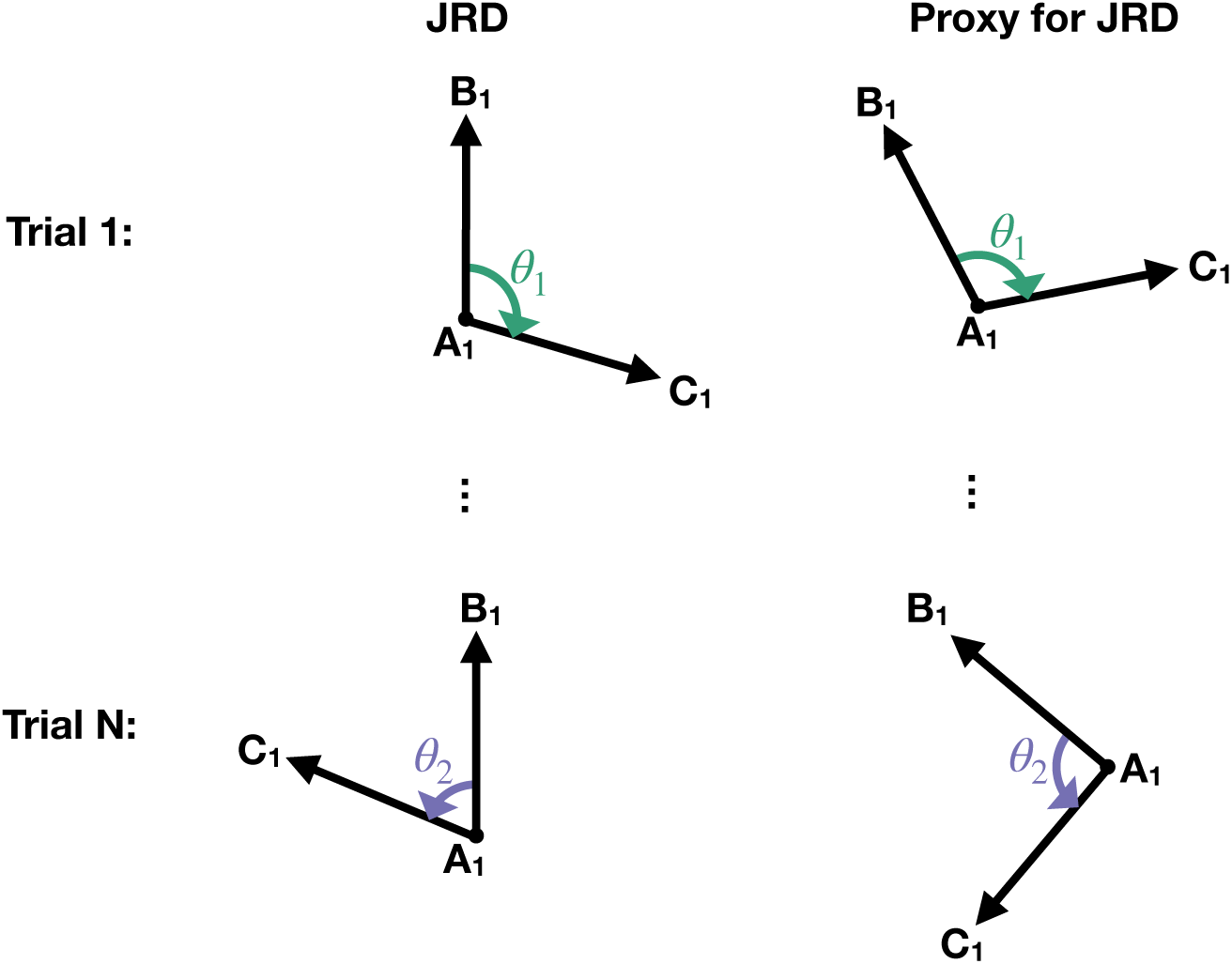
We developed a novel analytical technique to compare performance between the JRD task vs. the app-based in situ Direction Estimation Task and the Map Drawing Task. In the JRD task, we asked participants to imagine that they are standing at one location, facing a second location, and then to point to a third location (“Imagine you are standing at A facing B. Point to C.” Here, we can transform the app-based pointing and Map Drawing estimates into a JRD-like metric by taking the difference between the angles on two app-based pointing trials or two angles from the Map Drawing Task. Specifically, we can calculate the angle at which the participant pointed on the two trials that make up each of the angles in the JRD task (i.e., the first angle would be the angle they pointed for the B location while standing at A and the second angle would be the angle they pointed for the C location while standing at A). For the app-based in situ Direction Estimation Task, we then calculated the errors for our JRD-like metric vs. the actual angle, and we compared 1) correlations of the average performance on both tasks across participants (see Figure 4) and 2) the partial correlation between the patterns of errors on each trial (as a measure of the correlation in patterns of errors; note: we also performed this same procedure to compare the JRD task with the Map-Drawing Task; see Figure 5).

#### 2.2.2. Session Two: App-Based Navigation and in situ Direction Estimation Task

Participants completed Session Two on the Colby College campus using our custom-built iPhone application (which we developed in Swift/Xcode). Session Two lasted approximately 1.5 hours. Participants began by installing and launching the application (both TestFlight and our custom app), and then entered their participant number. Participants were asked to enable their location services and orientation sensors for the duration of the task.

Participants then reviewed on-screen instructions and safety guidelines. Participants then input the list of their 10 most familiar campus landmarks (which we emailed to them following the first session; see 2.2.1.). For the remainder of the app experiment, participants performed a Navigation Task, which was interspersed with our in situ Direction Estimation Task (see below). After completing all tasks, participants recorded their end time and exported the data from the application and to the research team. Participants then completed a post-experiment online questionnaire (we did not analyze the questionnaire here).

##### App-based Navigation Task

In the Navigation Task, participants walked to each of the 10 familiar landmarks identified in Session One. We randomly shuffled the order of the 10 landmarks before the start of the task. Upon arriving at each location, they indicated arrival within the application by pressing a button that said, “I have arrived.” We defined these arrival coordinates as the “ground truth” locations for each landmark for all of our analyses of spatial performance. At each location, participants completed our in situ Direction Estimation judgments (see below) for each of the remaining nine locations on their list.

##### App-based in situ Direction Estimation Task

In the in situ Direction Estimation Task, we asked participants to hold their phone parallel to the ground and point in the straight-line direction of a specified landmark. Each in situ Direction Estimation trial was immediately followed by a Confidence Rating, in which participants rated their confidence in the accuracy of their directional judgment on a 6-point Likert scale ranging from 1 (low confidence) to 6 (high confidence; note that we did not analyze the confidence data here). We repeated this procedure for each of the 9 other locations at each travel location, yielding direction estimates and confidence ratings for all ordered pairs of locations (here, there were 10 unique standing locations [determined by the Navigation Task] and 9 other locations for each standing location, resulting in 90 trials total).

We quantified accuracy on the in situ Direction Estimation Task using absolute angular error, defined as the absolute difference between the participant’s response angle and the true bearing to the target landmark. For each trial, the app recorded the participant’s estimated angle (“Response”) using the compass from the participants iPhone. We calculated the correct angle (“Answer”) by taking the coordinates from each of the “arrival” locations from the navigation task as the true landmark locations. We then calculated the ground truth (“Answer”) angles using the function `calculate_bearing` from OSMnx in Python (Boeing, 2017). As in the JRD Task, we used the median absolute angular error as the primary summary statistic for each participant to reduce the influence of outliers (i.e., the median is less influenced by outliers than the mean and here we had a relatively large number of trials, 90).

We were also interested in determining whether alternative measures of the app-based in situ pointing would be more strongly related to the lab-based tasks. Specifically, we developed a JRD proxy based on the app-based pointing data by gathering the “heading” and “pointing” response angles from the app task for each JRD question. We then calculated the angular difference between these angles to create our app-predicted JRD pointing response, as in our analysis of the map data (see 2.2.1 above, Figure 1, and Huffman et al., 2026; Huffman & Ekstrom, 2019). We analyzed the app-predicted pointing responses in two separate analyses. First, we report the correlation between the median app-predicted JRD pointing errors and the lab-based measures (similar to above, except using our app-predicted JRD pointing errors). Here, we calculated the median angular errors between the app-predicted JRD responses and the answers using the same method as we used for the JRD task. Second, we report the partial circular correlation, a measure of correlated errors, between the app-predicted JRD pointing responses and the lab-based measures, while taking into account the actual answers^1^ (e.g., as in Huffman et al., 2026; also see 2.3.2. below).

### 2.3. Analytical Methods

#### 2.3.1. Bayesian Methods

We were interested in determining whether or not there was a significant relationship between tasks (i.e., the null was a meaningful and interesting test because it would suggest participants used largely dissociable underlying representations to solve the tasks); thus, we employed a fully Bayesian framework. Specifically, for correlations, we calculated the Pearson correlation via the `correlationBF` function from the BayesFactor package (Morey & Rouder, 2012) with the default r-scale values (`rscale=“medium”,` which is √(2)/2). Likewise, for the t-tests (first, a paired t-test to compare the magnitude of errors on the app-based Direction Estimation Task and the JRD Task; second, to compare if the distribution of group Pearson r-to-z transformed partial correlation values were greater than zero; third, to compare if the map drawing data were better fit by the affine vs. Euclidean model using the ΔAIC across participants), we used the `ttestBF` function from the BayesFactor package (Rouder et al., 2009), again with the default r-scale value (`rscale=“medium”`, which is 1/3).

#### 2.3.2. Partial Correlation Methods

To investigate whether the within-participant patterns of errors were correlated between tasks, we implemented a partial correlation approach. For each trial of the JRD task, we also generated an app-based prediction of JRD performance and a Map-Drawing-based prediction of JRD performance using the participants’ angular estimates for both the “facing” and “pointing” trials within each of these tasks (for a similar approach with the JRD Task and the Map Drawing Task see: Huffman et al., 2026). We then calculated partial circular correlation coefficients using the `cor.circular` function from the circular package in R and custom-written code that implemented the following equation (Krzanowski, 2000):

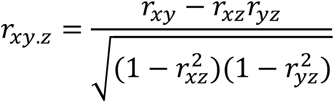

where *x* represents the circular array of angular responses based on one of the task conditions of interest and *y* represents another condition (we compared all possible combinations of all 3 of our tasks: JRD, Map-Drawing-based JRD prediction, app-based JRD prediction), *z* represents the circular array of angular answers for each of the questions on the JRD Task, and *r* represents the circular correlation coefficient for a given combination.

#### 2.3.1. Participant Exclusion Methods

To determine if each participant performed better than chance on the JRD Task and the in situ Direction Estimation Task, we used a permutation procedure, which we based on the procedure for the JRD Task in previous work (Huffman & Ekstrom, 2019). Specifically, we randomly shuffled the data and calculated the median angular error for each permutation. We performed this procedure 10,000 times to generate a null distribution of median error values. We then used a one-tailed test (here, to potentially allow more participants to be usedin our analysis; as in Huffman & Ekstrom, 2019) to determine whether each participant’s empirical median angular error was within the bottom 5% of their permutation-derived null distribution (i.e., a permutation-based, one-tailed p < .05; Ernst, 2004):

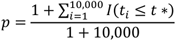

where *p* indicates the permutation-based p-value, *I()* is the indicator function that returns 1 if the condition is true and zero otherwise, *t_i_* indicates the test statistic for permutation *i* (here, the median error), and *t\** indicates the empirically observed test statistic (here, the median error for a given participant).

We also used the map drawing performance as an exclusion criterion: if the Euclidean model did not provide a better fit of a participant’s map drawing than a null model (we set significance as a ΔAIC value less than −2), then we flagged the participant for exclusion.

Altogether, we set an inclusion threshold that participants had to perform significantly better than chance (as defined above) on the JRD Task, the in situ Direction Estimation Task, and the Map Drawing Task to be included in our final analysis (i.e., if a participant did not perform significantly better than chance on one or more tasks, then we excluded them from our final analysis). We plot performance showing the excluded participants in Supplemental Figure 1. For further rationale on participant exclusion, please see above (section 2.1.).

### 2.4. Materials Availability Statement

We will make all analysis code available via github upon publication. We cannot share the raw data due to privacy concerns, but we will share minimally preprocessed data upon publication.

## 3. Results

### 3.1. Spatial Task Performance Differs Systematically Across Modalities

Participants exhibited distinct performance profiles across the three spatial tasks (Figure 2), indicating that task modality shaped both the mean performance and inter-individual variability. Across the two pointing tasks, performance differed in both accuracy and variability. App-based pointing yielded a lower median error (median = 11.42°, IQR = 4.70°) than the JRD task (median = 27.63°, IQR = 13.14°), which was also supported by paired t-tests of subject-level median errors (*t(57) =* 10.6, *p <* .001, BF_10_ = 1.65 × 10^12^). Together, these results indicate that abstract directional judgments in the JRD task produced higher and more variable errors across participants compared with app-based pointing.

**Figure 2:**
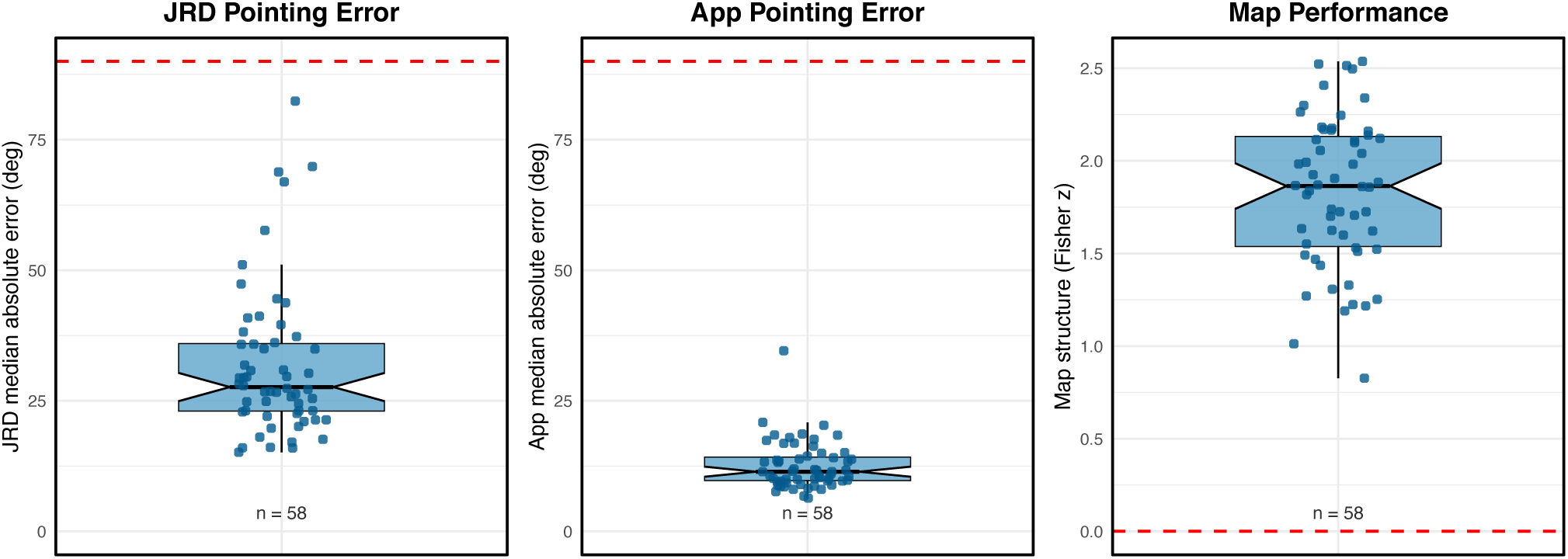
Performance differed systematically across spatial navigation tasks, with lower angular error for app-based pointing than JRD pointing and consistently above-chance structure in map-based reconstruction. Boxplots depict participant-level performance across three spatial navigation tasks: (A) judgments of relative direction (JRD), (B) app-based pointing, and (C) map-based reconstruction. JRD and App performance are expressed as subject-level median absolute angular error (degrees), computed from raw response–answer differences. Map performance is expressed as a Fisher z–transformed Euclidean model-fit metric. Individual points represent participants (n = 58). Notches indicate approximate 95% confidence intervals around the median. Red dashed reference lines indicate chance-level error (90°) for the pointing tasks and zero structure for the map task.

Map-based performance showed consistently positive Fisher r-to-z values (median = 1.55, IQR = 0.483), indicating meaningful spatial structure overall while still revealing substantial individual differences in how well participants preserved metric relationships among landmarks (see Supplemental Figure 1C for data across all participants, including excluded participants).

### 3.2. We Observed a Correlation between Real-World Memory Performance on the Lab-Based JRD Task and Map Drawing Task

We next examined the relationship between the two laboratory-based measures of real-world spatial memory: errors on the JRD Task and model fit in the Map Drawing Task. We found a negative correlation between the JRD median absolute error and the Map Drawing Task fit score (r = −.50), with strong Bayesian evidence for a nonzero relationship (BF₁₀ = 428.91; see Figure 3A). Participants who showed lower directional error on the JRD task tended to produce more accurate maps, as indexed by higher Fisher z–transformed bidimensional regression fits. These results support previous work that has compared performance between the JRD Task and the Map Drawing Task for both virtual environments and large-scale, real-world environments (e.g, Huffman et al., 2026; Huffman & Ekstrom, 2019), thus suggesting that these tasks tap into partially overlapping cognitive representations.

**Figure 3:**
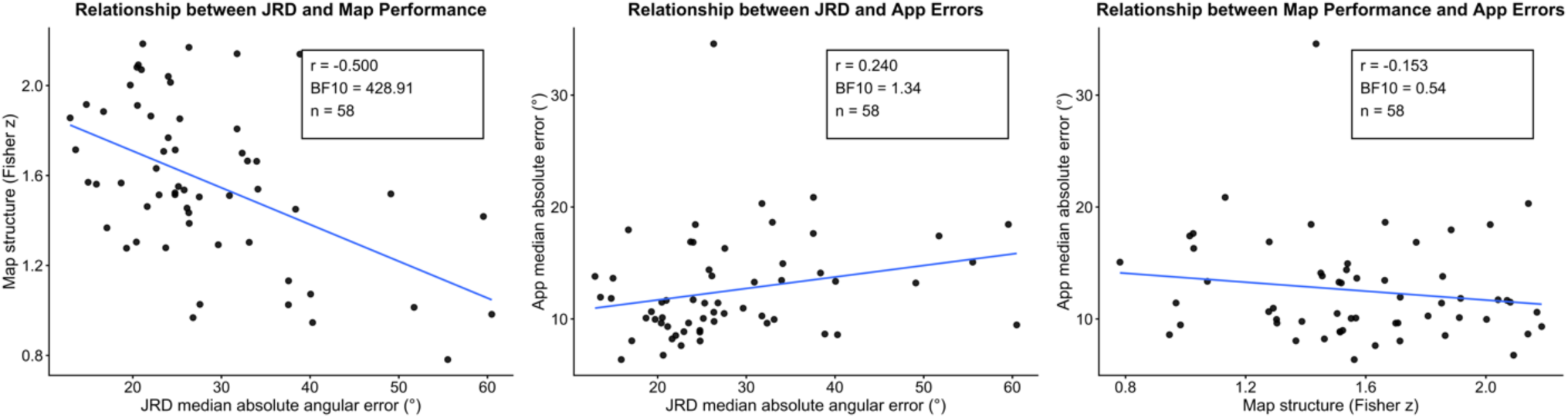
We observed a strong correlation between lab-based measures but inconclusive correlations between the lab-based measures and the in situ App pointing accuracy: (A) We observed strong evidence of a correlation between JRD median absolute error and Map Drawing performance, (B) We observed very weak evidence for a correlation between JRD median absolute error and in situ app-based pointing median absolute error, and (C) We observed very weak evidence in favor of a null correlation between in situ app-based pointing median absolute error and Map Drawing performance. Solid lines indicate least-squares regression fits. Insets report Pearson’s correlation coefficient (r), Bayes factor (BF_10_), and sample size (n = 58). JRD and app-based median pointing errors are expressed in degrees; map structure is expressed as the Fisher z–transformed bidimensional fit.

### 3.3. The Raw App-Based in situ Pointing Error did not Strongly Correlate with Either the Lab-Based JRD Task or Map Drawing Task

We next examined how app-based pointing performance related to the two laboratory measures. We found that there was only a very weak positive correlation between median error on the JRD Task and the app-based in situ Direction Estimation Task error (r = .24) and the Bayes factor for this relationship was inconclusive (BF_10_ = 1.34; see Figure 3B), indicating inconclusive support for a possible association between the two pointing measures (i.e., neither in favor of the alternative nor the null).

We next compared the correlation between the app-based in situ Direction Estimation Task and the Map Drawing task, and we observed an even weaker correlation (r = −.15) and the Bayes factor was inconclusive and only very weakly favored the null hypothesis (BF_10_= 0.54; see Figure 3C).

Taken together, while the Bayes factors are inconclusive, these results suggest a lack of a strong association between lab-based measures of spatial performance and the raw app-based in situ Direction Estimation task performance (e.g., perhaps a seeming dissociation between tasks). While JRD performance showed a strong relationship with Map Drawing Performance, app-based pointing did not exhibit strong correlations with either lab-based measure, thus suggesting that the raw app-based pointing accuracy and traditional laboratory tasks do not capture identical sources of variance in spatial memory. However, we note that the Bayes factors were inconclusive so we do not want to overinterpret our findings here and, moreover, we found a very different pattern of results in subsequent analyses in which we generated an app-based prediction of angular judgments between multiple landmarks simultaneously (see 3.4.).

### 3.4. We Found Significant Correlations between an App-based Prediction of JRD Performance and both the Lab-Based JRD Task and the Map Drawing Task

We aimed to determine whether alternative measures of the app-based pointing would be more strongly related to the lab-based tasks. Specifically, one notable difference between the app-based pointing vs. the JRD and map drawing tasks is that the app-based pointing task only consists of pointing to a single landmark from a given standing location, whereas the other tasks are inherently relational and comprise estimates from more than two landmarks at a time. Thus, we were interested in developing a JRD proxy based on the app-based pointing data (i.e., as in our analysis patterns of errors on the JRD Task and the Map Drawing Task below; for the logic of our analysis see 2.2.2. and Figure 1; Huffman et al., 2026; Huffman & Ekstrom, 2019).

We found that the app-predicted JRD pointing errors were strongly positively correlated with JRD pointing errors (r = .58, BF₁₀ = 1.12 × 10⁴, n = 58; see Figure 4A) and negatively correlated with Map Drawing Task accuracy (r = −.45, BF₁₀ = 85.89, n = 58; see Figure 4B). These findings indicate that expressing app-based pointing performance within the same angular reference frame as the JRD reveals strong correlations between tasks. More broadly, these results suggest that the previously weak evidence for relationships between app-based pointing and laboratory measures at least partially reflects differences in task measures (e.g., relational vs. isolated) and structure rather than an absence of a shared spatial representation.

**Figure 4:**
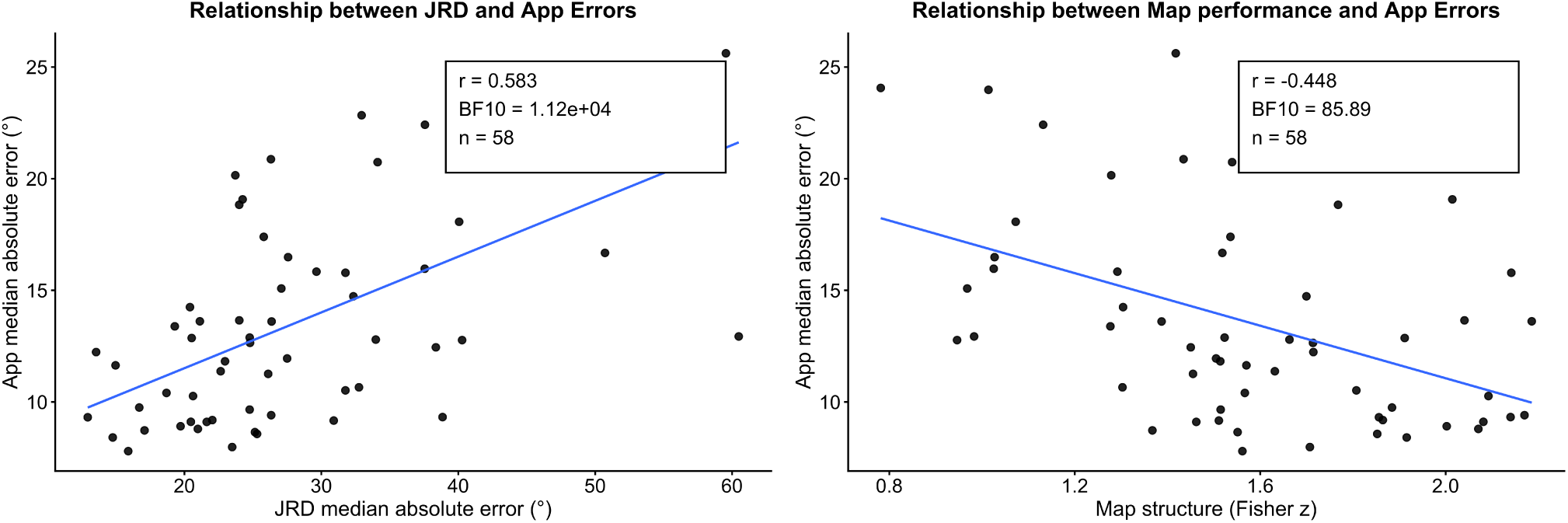
We found strong correlations between the predicted JRD responses from app-based pointing error and both the median absolute error on the JRD Task (left panel) and performance on the Map Drawing Task (right panel). Each point represents one participant. The solid line indicates the least-squares regression fit. Reported values reflect Pearson’s correlation coefficient (r), sample size (n), and Bayes factor (BF_10_) quantifying evidence for a nonzero association.

### 3.5. A Partial Circular Correlation Analysis Revealed that the Patterns of Errors across the App-based Pointing, the JRD Task, and the Map Drawing Task are all Correlated

We next report the results of a partial correlation analyses to examine whether the patterns of errors were shared across tasks shared across measures. Here, we aimed to determine whether the patterns of errors are correlated between tasks (Huffman et al., 2026; Huffman & Ekstrom, 2019) by generating predictions of JRD performance across both the map drawing data and the app pointing data (see 2.3.2.). If participants employ similar underlying cognitive representations to solve the tasks, then we should observe partial correlations that are greater than 0 (i.e., indicating stable and consistent correlations in patterns of errors across tasks). We found positive mean partial correlations across all three task pairs, with the Bayes factors strongly favoring the alternative hypothesis in each case, indicating reliable associations in directional judgments between all tasks. Specifically, we found a positive partial correlation between the two lab-based tasks: JRD performance and Map-Drawing-predicted JRD (mean z[r] partial correlation = 0.47; BF₁₀ = 1.23 × 10¹⁶; see Figure 5), thus replicating previous work (Huffman et al., 2026). Moreover, we observed evidence of positively correlated errors between the lab-based and app-based measures: the partial correlations between the app-predicted JRD and the empirical JRD performance (mean z[r] partial correlation = 0.29; BF₁₀ = 2.41 × 10¹¹; see Figure 5) and the app-predicted JRD and the Map-Drawing-predicted JRD (mean z[r] partial correlation = 0.43; BF₁₀ = 1.20 × 10¹⁵; see Figure 5) both exhibited extreme evidence in favor of the hypothesis that the partial correlations are greater than 0. Therefore, our results strongly suggest that app-based pointing, JRD performance, and map drawing all share partially overlapping patterns of error structure that cannot be explained solely by shared variance with the answers.

**Figure 5:**
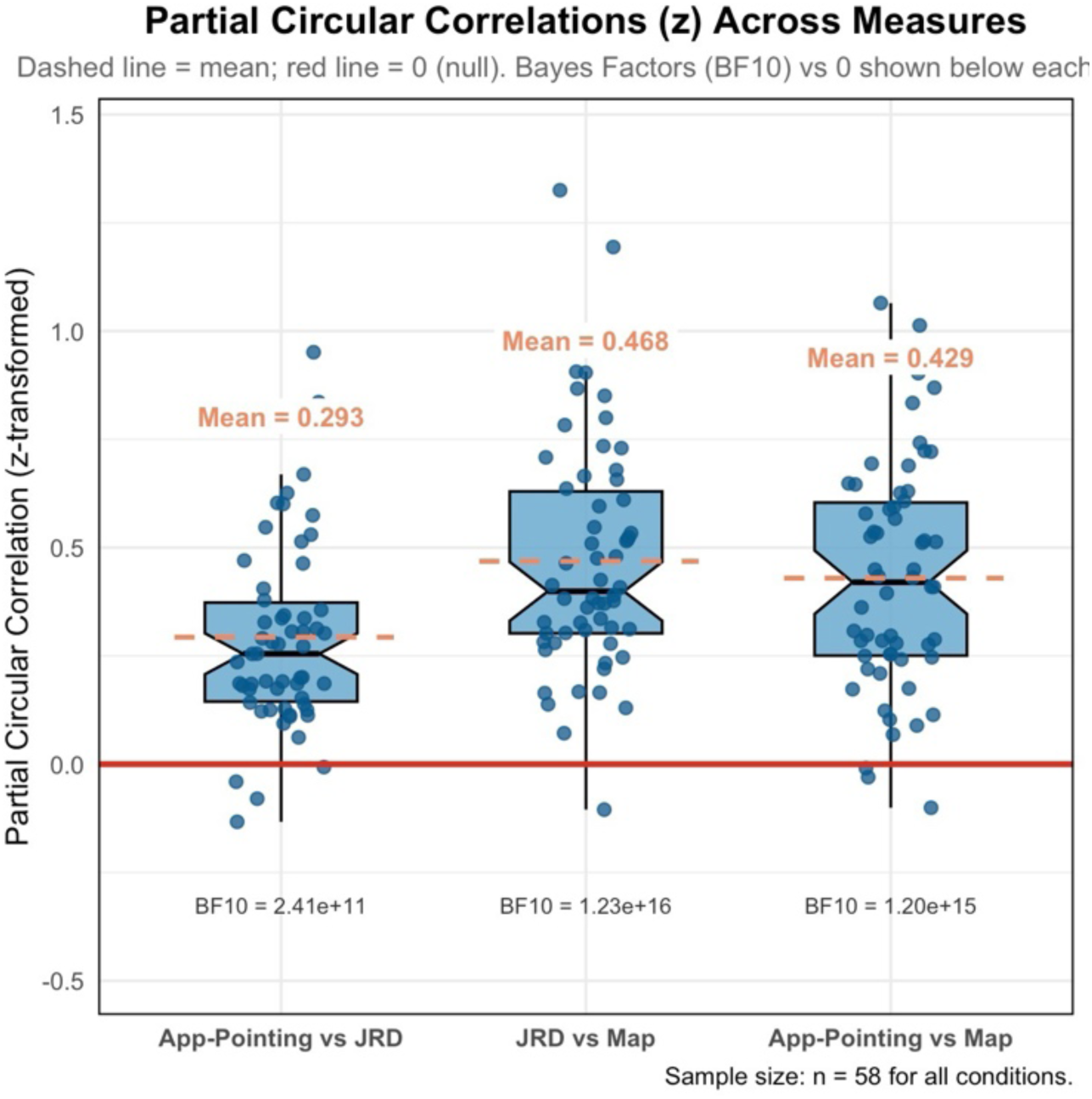
Evidence that all of the tasks exhibit correlated errors. Here, the boxplots show the Fisher z–transformed partial circular correlations between directional judgments across tasks while partialling out the answers: app-based pointing versus JRD (left), JRD versus Map Drawing (center), and app-based pointing versus Map Drawing (right). Individual points represent participant-level partial correlations. Dashed lines indicate mean values, and the solid red line indicates the null value (z = 0). Bayes factors (BF₁₀) comparing each distribution against zero are shown below each box. Sample size was n = 58 for all task pairs. For more information about our approach, see Figure 3.

Altogether, we found strong correlations between performance of the app-predicted JRD performance with lab-based measures as well as strong evidence of partial correlations in error patterns between all of the tasks. Thus, our findings suggest that, when we transform the app-based pointing data into a more comparable framework to the lab-based tasks, we observe strong evidence for relationships between the tasks. These findings represent an important advancement by showing that our novel analytical technique uncovers partially overlapping cognitive representations between all 3 tasks. We also note that these results stand in stark contrast with the comparison of the raw app-based in situ Direction Estimation Task with the lab-based measures (see 3.3.). We discuss these findings in more detail in the Discussion.

### 3.6. The Map Drawing Data were Better Fit by an Affine Model, thus Providing Evidence for Systematic Distortions in Spatial Memory

We next tested whether the participants’ memories exhibited systematic distortions by comparing the fit of the Map Drawing Task data by an affine model, which allows for systematic distortions (including shearing; e.g., accommodating alignment and rotation heuristics; Nakaya, 1997), vs. a Euclidean model (i.e., translation, rotation, and uniform scaling). A Bayesian one-sample *t*-test comparing ΔAIC values against zero (see Supplemental Methods) yielded extreme evidence for the affine model over the Euclidean model (BF₁₀ = 1,158.83), and the median ΔAIC reached the conventional threshold for significant model preference, thus suggesting a significant effect, on average, within individual participants (median = −2.38; see Supplemental Figure 2; for more details see Supplemental Methods and Supplemental Results; Huffman et al., 2026; Nakaya, 1997). Altogether, these results suggest that participants’ memories for a familiar, personally relevant, large-scale, real-world environment are systematically distorted (thus replicating Huffman et al., 2026).

## 4. Discussion

We applied several analyses to test the correlation between real-world memory performance with our app-based tasks and lab-based tasks for a large-scale, familiar environment: the college campus. We found a strong correlation between performance on our lab-based tasks—the JRD Task and the Map Drawing Task—participants that pointed more accurately tended to also draw more accurate maps. We also found strong evidence for a positive partial correlation between JRD pointing responses and a Map-Task prediction of JRD pointing responses, which suggests that the patterns of errors were correlated between tasks. Altogether, these findings suggest that participants use partially overlapping cognitive representations to solve the lab-based tasks. Our findings replicate previous work that examined new learning within a virtual environment (Huffman & Ekstrom, 2019) as well as a recent study that extended these findings to real-world environments (Huffman et al., 2026). Note that in contrast to our findings, Bryant (1984) found a much more modest relationship between patterns of errors analyzed at the group level, which they argued challenges the construct validity of these tasks. We note that our approach here is different in that we analyzed the data within each participant separately, thus potentially opening a more sensitive measure with greater construct validity (i.e., we argue it is preferable to study spatial memory patterns at the level of individual participants instead of averaging between participants, especially for measuring patterns of errors). We think that it will be interesting for future studies to further investigate the conditions under which patterns of errors are correlated vs. uncorrelated between the JRD task and the Map Drawing Task. Altogether, our results from the lab tasks further support the construct validity of these measures and suggest that they tap into holistic, abstract representations of the environment.

Our results add to a growing literature investigating real-world spatial memory in lab vs. in situ and offer a key advancement to studying memory for real-world environments. Our main goal was to develop, validate, and compare performance of a novel app-based measure of spatial memory vs. the lab-based tasks. We found evidence for both shared and unique variance between our lab and in situ memory tasks. We found minimal evidence for a correlation between the raw app-based in situ Direction Estimation Task performance and the lab-based tasks. If taken at face value, these results suggest that the in situ Direction Estimation Task and the lab tasks tap into largely dissociable cognitive representations, which could support previous work that has shown a dissociation between in situ pointing vs. JRD pointing (e.g., Waller & Hodgson, 2006; Zhang et al., 2014; for similar ideas with respect to in situ pointing with a manipulation of disorientation see Wang & Spelke, 2000, 2002).

We were also interested in determining whether the patterns of pointing responses were correlated between the tasks by calculating the predicted JRD angles by comparing the pairs angles between the two corresponding trials for the heading and pointing judgments of the in situ Direction Estimation Task. Here, we found strong evidence of a correlation between the app-based in situ Direction Estimation Task and the lab tasks, based both on the performance across participants as well as the putative measure of correlated errors (i.e., the partial correlation analysis). Altogether, these results suggest that when we convert the in situ pointing responses into a common framework with the lab tasks, we see much clearer evidence of a relationship between tasks. Moreover, the finding of significant partial correlations provides stronger evidence that the patterns of errors were correlated, thus suggesting these tasks tap into partially overlapping cognitive representations. Therefore, our findings are a key extension on previous findings that focused on real-world memory using lab-based measures (Huffman et al., 2026), and we think it will be interesting for future research to employ our methods to further uncover conditions in which performance and patterns of errors are correlated vs. uncorrelated for the lab and in situ spatial memory tasks. Moreover, our findings extend previous research that has shown correlations between new learning of desktop-based virtual environments and real-world environments (Coutrot et al., 2019; Puthusseryppady et al., 2022) into the realm of memory for familiar, personally meaningful, large-scale spaces that are learned over the course of months to years. Thus, our findings lay an important foundation for future studies in different populations.

Disruptions in navigation are among the earliest neurocognitive changes in Alzheimer’s disease (AD), potentially emerging years or even decades before impairments are detected by standard neuropsychological tests (Allison et al., 2016; Coughlan et al., 2018; Vlček & Laczó, 2014), and there may be a dissociation between getting disoriented in familiar environments, such that people at risk of AD become disoriented whereas healthy older adults do not (Rosenbaum et al., 2012). Additionally, the brain network that supports spatial navigation, including the hippocampus, entorhinal cortex, and retrosplenial cortex, are also regions that are particularly vulnerable to early pathological change, thus theoretically making navigation a sensitive and specific window into presymptomatic AD (Braak & Del Tredici, 2015; Coughlan et al., 2018, 2019; Kunz et al., 2015; Moffat, 2009; Puthusseryppady et al., 2022; Vlček & Laczó, 2014); however, to date, the implementation of spatial navigation within large-scale, real-world, familiar environments as an early biomarker of cognitive decline is limited, due in large part to technological limitations (e.g., limited means of studying participants within their own large-scale environments in which they are living; for similar arguments about the lack of research on the connection between real-world spatial memory vs. virtual environments in AD see: Puthusseryppady et al., 2020), thus our methods and findings here provide an important first step toward future studies in different populations, especially people at risk of developing AD.

We found that real-world memory performance was better and less variable on the in situ Direction Estimation Task than the JRD Task, which we think will be important for future work that is aimed at developing tasks for detecting preclinical changes in spatial memory in people at risk of developing AD. Our findings here build on previous studies that have reported better pointing accuracy for tasks in which the participants have access to visual information about their heading direction in the environment (including pictures or views of virtual environments during pointing) vs. the JRD task in which participants need to imagine both the heading and the direction of the target (e.g., Stevens & Carlson, 2016; Zhang et al., 2014). Moreover, our findings are an important extension because we directly measured performance within individuals both as they were situated in the environment vs. the JRD task. We also note that previous studies have reported a large range of individual variability in performance on spatial memory tasks, especially those that tap into abstract measures of memory, such as the JRD Task and the Map Drawing Task (e.g., many otherwise healthy individuals can perform poorly or even at chance levels even after repeated exposure to the environment; e.g., Ishikawa & Montello, 2006; Weisberg et al., 2014; Weisberg & Newcombe, 2018; Wolbers & Hegarty, 2010). As an important extension, we think it is interesting that we found strong correlations when we converted the app-based pointing into a JRD-like metric, both via the correlations across participants and the correlated patterns of errors, especially given the better performance overall for the raw app-based pointing. In summary, in the context of developing sensitive and specific measures for early neurocognitive changes in presymptomatic AD, we argue that the baseline of better performance with less interindividual variability on the in situ spatial memory task could allow for more diagnostic measures, including tracking longitudinal changes in performance.

Recent theories have argued that there is a possible disconnect between studying memory in the lab (e.g., with virtual environments) and the real world (Puthusseryppady et al., 2020; Taube et al., 2013). Furthermore, the majority of studies of changes in spatial memory in aging and AD have focused on new spatial learning, but we hypothesize that studying memory for familiar spatial environments would more closely mimic the real-life differences between healthy aging and AD. For example, patients with AD often get disoriented and lost within familiar spatial environments (Coughlan et al., 2018; Puthusseryppady et al., 2022; Vlček & Laczó, 2014), which is rare for healthy older adults. In fact, the limited work on memory for familiar environments in healthy older adults suggests that they may not have large changes in memory performance for familiar environments relative to young adults (Rosenbaum et al., 2012), which stands in stark contrast to the near consensus for age-related impairments in new spatial learning relative to young adults (e.g., Lester et al., 2017; Moffat et al., 2001; Moffat & Resnick, 2002; Spiers et al., 2023), especially within virtual environments. Moreover, previous work has suggested that memory differences in healthy aging can be mitigated under more immersive conditions vs. desktop virtual environments (Hill et al., 2024; for related discussion of the role of body-based cues in aging research see: Diersch & Wolbers, 2019). Altogether, we posit that real-world spatial memory, especially within familiar environments, will have higher sensitivity and specificity for detecting preclinical signs of AD (i.e., in differentiating healthy vs. preclinical pathological aging). In conclusion, we found evidence of partially overlapping representations for our lab-based measures as well as evidence of both unique and shared variance, which varied as a function of methodological approach, between our lab-vs. real-world-based measures of spatial memory. Our findings provide an important theoretical and methodological advancement in terms of understanding human spatial memory for large-scale, real-world, familiar environments that are learned over the course of months to years. Additionally, we argue that studying spatial memory for familiar, large-scale, real-world environments could provide a diagnostic tool for distinguishing between healthy-aging and preclinical AD, and our findings here take an important first step in that direction by employing a set of new tools within healthy young adults.

## Generative AI Disclosure Statement

We did not use generative AI (e.g., LLMs) in writing this manuscript (i.e., we wrote everything ourselves).

## Supplemental Material

### Supplemental Methods

#### Supplemental Analysis of the Map Drawing Task Performance

To assess whether participants’ map-based reconstructions exhibited systematic spatial distortions, we compared the fit of a so-called Euclidean transformation and an affine transformation (allowing for non-uniform scaling and shearing) for each participant’s map using bidimensional regression (Carbon, 2013; Friedman & Kohler, 2003; Tobler, 1994). We conducted model comparison using differences in Akaike Information Criterion (ΔAIC = AIC_affine − AIC_Euclidean), with negative values indicating stronger support for the affine model and positive values indicating stronger support for the Euclidean model, with |ΔAIC| > 2 often taken as evidence of a significant difference between models (Nakaya, 1997). We ran the model fit at the level of individual participants. We then aggregated the ΔAIC values across participants and compared whether these values differed from zero using a Bayesian one-sample *t*-test (see Supplemental Figure 2). Because model fits were evaluated separately for each participant, this analysis constitutes a random-effects assessment of map structure (for a related approach see: Huffman et al., 2026).

^1^ We note that our a priori analytical plan was to calculate the partial correlation between all tasks, as in previous work (Huffman et al., 2026; Huffman and Ekstrom, 2019). We report the analysis of median errors from the JRD-proxy score as a supporting analysis for our partial correlation analysis; however, we would like to emphasize that we are reporting our full set of analyses (i.e., this analysis is not part of a “fishing expedition”).

### Supplemental Figures

**Supplemental Figure 1:**
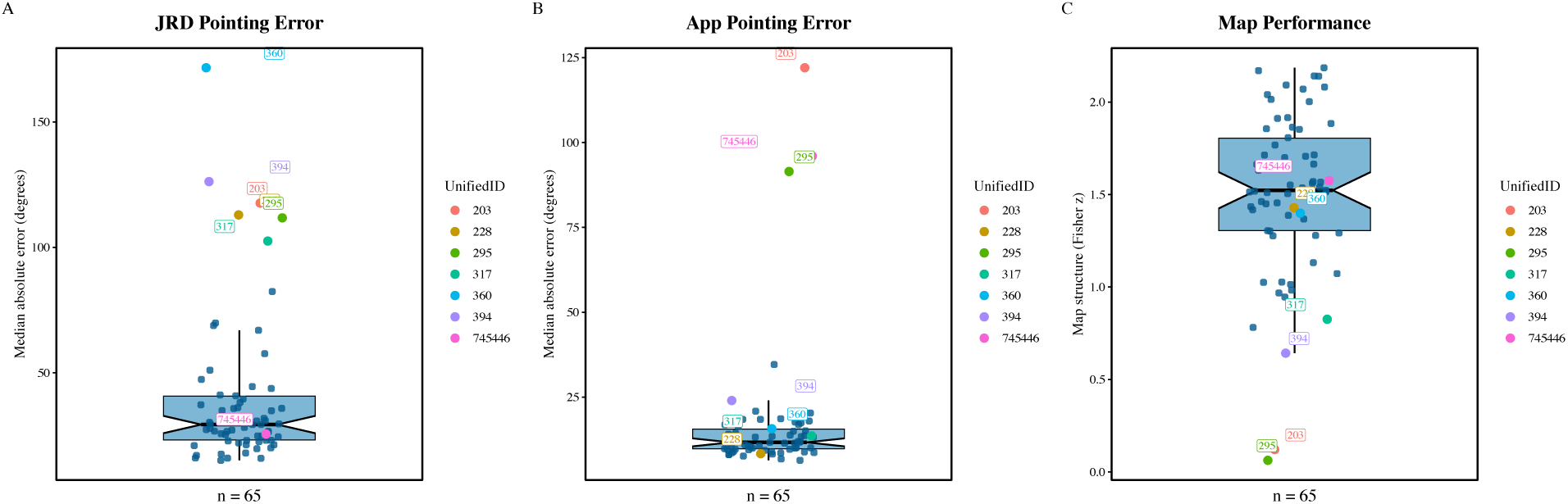
Boxplots summarize subject-level performance for the (A) computer-based JRD pointing error, (B) the app-based in situ Direction Estimation Task pointing error, and (C) Map Drawing performance, with jittered points showing individual participants (n = 65). For panels A–B, values reflect median absolute angular error (degrees); for panel C, values reflect map performance as the Fisher r-to-z transformed model fit values for the bidimensional regression. Colored, labeled points indicate participants flagged for exclusion based on the multimodal screening criteria.

**Supplemental Figure 2:**
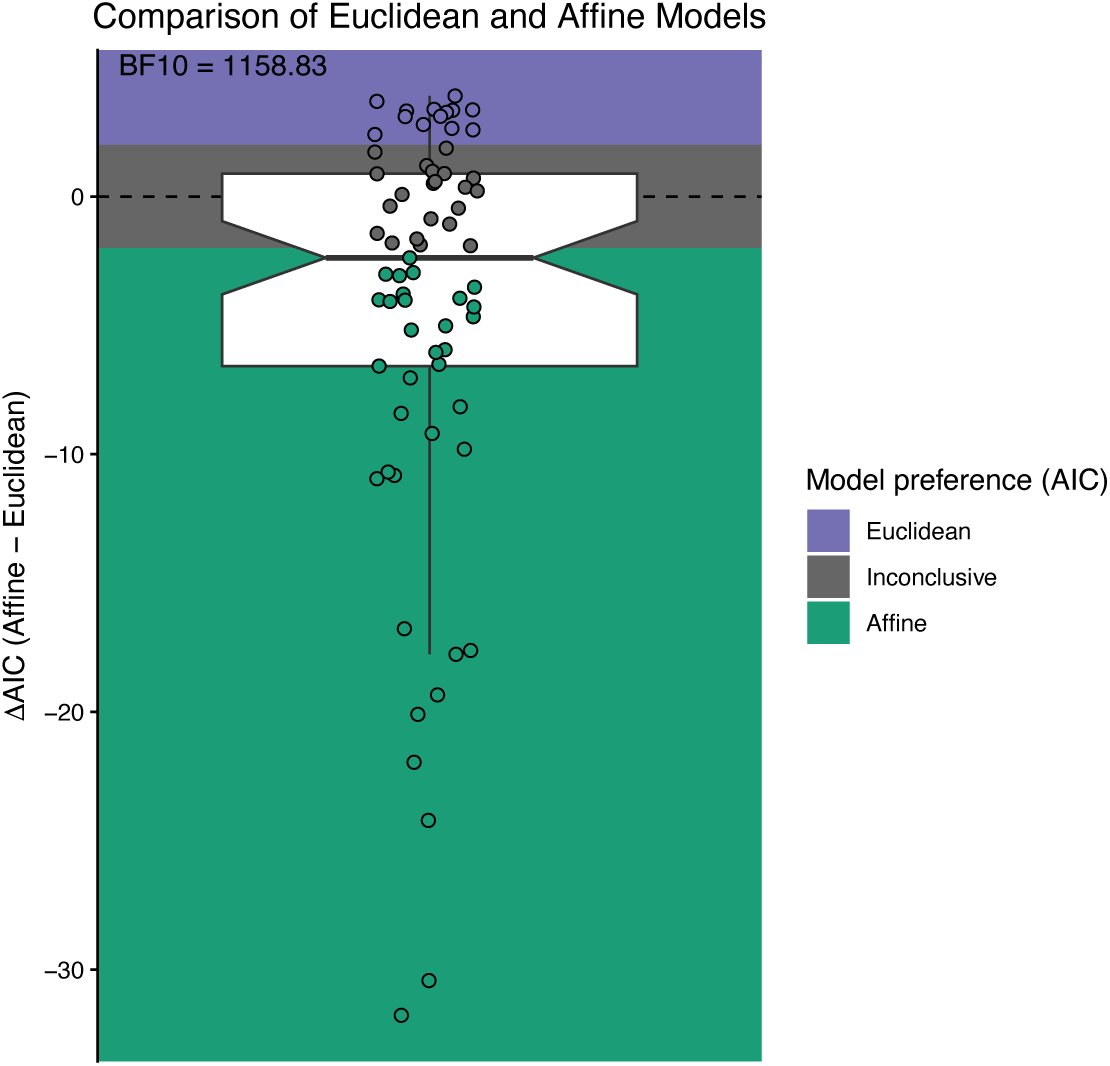
Evidence that participant’s maps exhibit systematic distortions (here, better fit by the affine than the Euclidean model). Each point represents an individual participant’s map-based reconstruction, plotted as the difference in Akaike Information Criterion between affine and Euclidean models (ΔAIC = AIC_affine − AIC_Euclidean). Positive values indicate stronger support for the Euclidean transformation (i.e., translation, rotation, and uniform scaling), whereas negative values indicate stronger support for an affine representation (i.e., translation, rotation, and non-uniform scaling), consistent with systematic distortions, such as alignment and rotation heuristics. Shaded regions denote model preference thresholds, with ΔAIC > 2 indicating significant evidence for the Euclidean model, ΔAIC < −2 indicating significant evidence for the affine model (at the level of individual participants), and intermediate values reflecting inconclusive preference. The dashed horizontal line marks ΔAIC = 0. The majority of participants were significantly better fit by the affine model and the Bayes factor t-test revealed that the average ΔAIC was less than 0. Each dot indicates the ΔAIC for an individual participant.

